# Insulin and Exercise-induced Phosphoproteomics of Human Skeletal Muscle Identify REPS1 as a New Regulator of Muscle Glucose Uptake

**DOI:** 10.1101/2023.11.10.566644

**Authors:** Jeppe Kjærgaard Larsen, Cecilie B. Lindqvist, Søren Jessen, Mario García-Ureña, Amy M. Ehrlich, Farina Schlabs, Júlia Prats Quesada, Johann H. Schmalbruch, Lewin Small, Martin Thomassen, Anders Krogh Lemminger, Kasper Eibye, Alba Gonzalez-Franquesa, Jacob V. Stidsen, Kurt Højlund, Tuomas O. Kilpeläinen, Jens Bangsbo, Jonas T. Treebak, Morten Hostrup, Atul S. Deshmukh

## Abstract

Skeletal muscle regulates glucose uptake in response to insulin and exercise which is critical for maintaining metabolic health. We conducted a comprehensive phosphoproteomic analysis of skeletal muscle from healthy people in response to an acute bout of exercise or insulin stimulation by a hyperinsulinemic euglycemic clamp. Our analysis revealed 233 phosphosites regulated by both exercise and insulin of which most phosphosites were regulated in opposite directions. However, 71 phosphosites on 55 proteins displayed regulation in the same direction, indicating a potential convergence of signaling pathways. We identified the vesicle-associated protein, REPS1, to be phosphorylated at Ser709 in response to both insulin and exercise. REPS1 protein level and Ser709 phosphorylation were closely related to insulin-stimulated glucose uptake in skeletal muscle and required for maximal insulin-stimulated glucose uptake. Furthermore, we observed that insulin triggered phosphorylation of REPS1 Ser709 via P90S6 kinase (RSK) and is impaired in mice and humans with insulin resistance. Collectively, REPS1 is a convergence point for insulin and exercise signaling and a promising therapeutic target in insulin resistance.

## Introduction

Skeletal muscle is the largest tissue in the human body and plays a pivotal role in maintaining glucose homeostasis and energy metabolism. One of its primary metabolic functions is the uptake and utilization of glucose, which is essential for normal physiological processes and is particularly relevant in the context of metabolic disorders, such as type 2 diabetes (T2D)^1^. Impaired insulin-stimulated skeletal muscle glucose uptake is a hallmark of insulin resistance and is a central feature of the pathogenesis of T2D^2^. Skeletal muscle glucose uptake is intricately regulated by multiple signaling pathways that dictate its delivery, transport, and metabolism^3–5^. Glucose transporter 4 (GLUT4) plays a central role in regulation of skeletal muscle glucose uptake^6,7^. During unstimulated conditions GLUT4 predominantly resides within intracellular vesicles. However, in response to insulin or exercise, GLUT4-containing vesicles translocate to the plasma membrane^5^. This suggests that exercise- and insulin-induced GLUT4 translocation rely on commonly regulated distal mechanisms and protein machinery.

Studies on human skeletal muscle in patients with T2D have revealed an impairment in insulin signaling pathways targeting GLUT4, while signaling induced by exercise towards GLUT4 translocation remains largely intact^8–10^. The study of insulin signaling has mainly been limited to two major pathways: phosphatidylinositol 3-kinase (PI3K)-AKT and RAC1-p21-activated kinase (PAK) pathways which are recognized as vital modulators of insulin-stimulated glucose^11–13^. In contrast, exercise-mediated glucose uptake is mainly orchestrated through distinct proximal signaling mechanisms integrating a spectrum of mechanical, chemical, and stress signals^14–18^. One hypothesis is that insulin and exercise-triggered signaling converge at distal nodes to meet common cellular needs e.g., an increase in glucose import. This notion was supported by the discovery of the molecular switches, TBC1D1 and TBC1D4, which intricately regulate the action of insulin and exercise on trafficking of GLUT4 and lipid transporter-containing vesicles to the cell surface^19–22^. Intriguingly, an acute bout of exercise can potentiate subsequent insulin signaling, notably exemplified by the AMPK-TBC1D4 axis in both human and rodent skeletal muscle^23–25^. Despite these few examples, our understanding of these converging distal mechanisms and the shared protein machinery remains incomplete and underexplored.

Recent improvements in mass spectrometry (MS)-based phosphoproteomics have advanced the study of dynamic protein signaling^26–28^. Pioneering studies in skeletal muscle have unveiled extensive signaling networks activated by both exercise and insulin^29–31^. Despite these valuable insights, the intricate interplay among these pathways and their regulation of GLUT4 remains to be fully elucidated. A comprehensive understanding of these interactions is crucial for unraveling therapeutic targets that can increase skeletal muscle glucose uptake in diseases characterized by insulin resistance. Based on the current evidence, we hypothesized that a core set of proteins, modulated by phosphorylation in response to both insulin and exercise, form a fundamental regulatory nexus for skeletal muscle glucose uptake.

We conducted a phosphoproteomic analysis in skeletal muscle from healthy individuals in response to an acute bout of exercise or insulin stimulation by a hyperinsulinemic euglycemic clamp. While the majority of phosphosites regulated by both insulin and exercise displayed opposite responses to the stimulus, we identified 71 sites on 55 proteins that were regulated in the same direction, potentially influencing glucose uptake. In particular, we found RalBP1-associated Eps domain-containing protein 1 (REPS1), a vesicle-associated protein, was phosphorylated in response to both insulin and exercise. We show that REPS1 is crucial for glucose uptake in skeletal muscle and that the phosphorylation depends on P90S6 kinase (RSK). Importantly, REPS1 S709 phosphorylation was impaired in response to insulin stimulation in several *in vivo* insulin resistant models and showed a strong correlation with insulin sensitivity. Our results position REPS1 at the intersection of insulin and exercise signaling and expand our understanding of the mechanisms influencing glucose uptake into skeletal muscle.

## Results

### Phosphoproteomic Signature of Insulin and Exercise Signaling in Human Skeletal Muscle

To accurately capture signaling events in response to insulin and exercise, we conducted two experimental trials, in randomized order, separated by 1-2 weeks on eight healthy males (age, 18–36 yrs., lean mass index, 14–22 kg/m2; maximal oxygen uptake [VO2max], 40–60 ml/kg/min). Given our objective to identify converging distal signaling events in response to insulin and exercise, we conducted both trials on the same individuals to minimize potential inter-subject variations in insulin and exercise signaling. During the experimental trials, participants underwent either a 2-h hyperinsulinemic euglycemic clamp (HEC) or performed a single bout of high-intensity cycling exercise for 10 min with maximal effort (Figure 1A). Vastus lateralis muscle biopsies were obtained immediately before and after the intervention on both experimental days. The acute bout of exercise reduced glycogen stores (*P* < 0.01), while no change was observed with insulin stimulation (Figure 1B). Insulin stimulation increased leg glucose uptake (*P* < 0.001, Figure 1C). We confirmed the activation of canonical insulin and exercise-induced signaling pathways in the muscle biopsies by Western blot analysis (Figure 1D). For instance, both insulin and acute exercise induced phosphorylation of mTOR at S2448, while insulin specifically increased p-AKT at S473 and exercise specifically increased p-AMPK at T172 and p-ACC at S221 (Figure 1D & Figure S1A-D).

**Figure 1.**
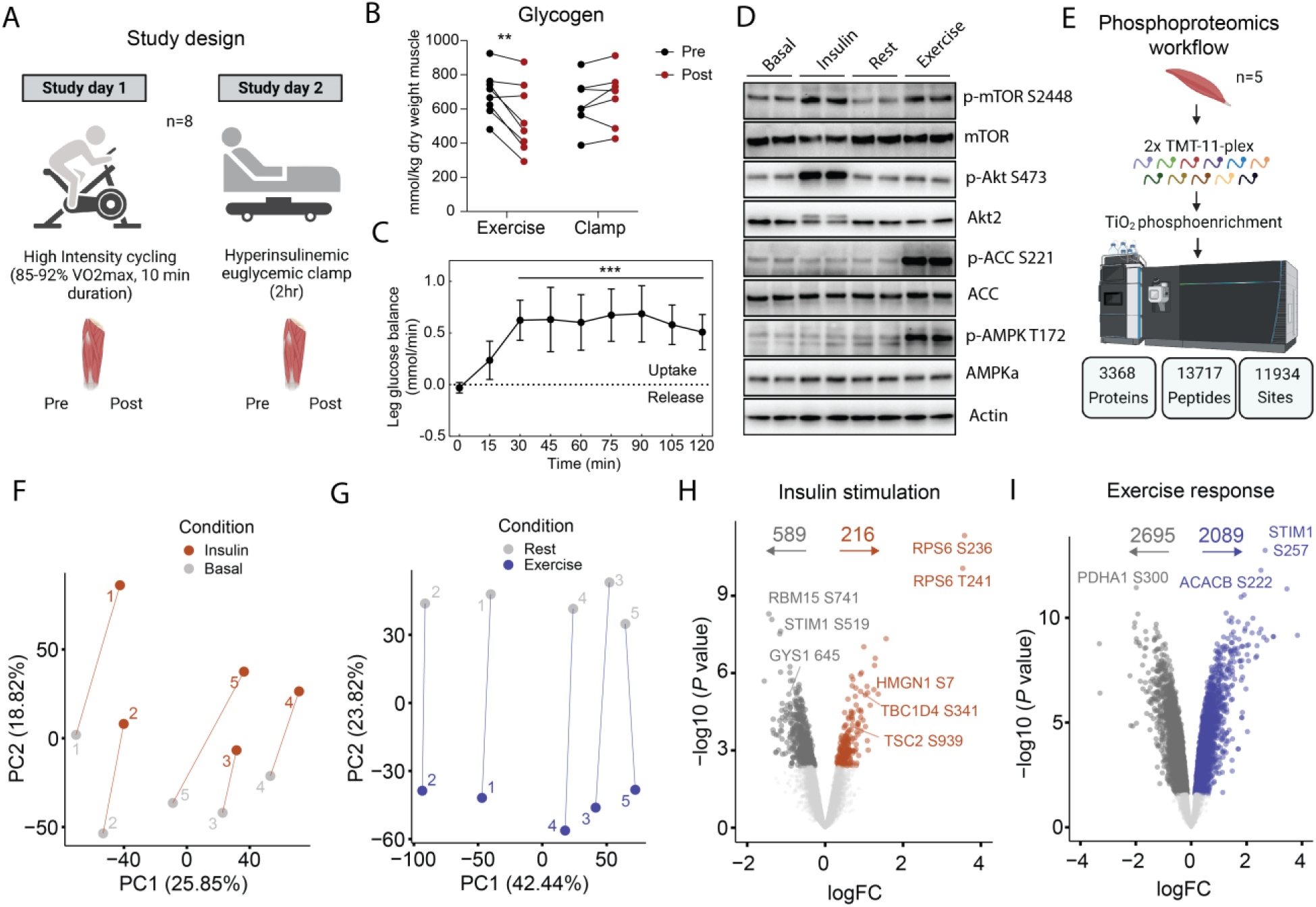
Phosphoproteomic Signature of Insulin and Exercise Signaling in Human Skeletal Muscle. Study design – Eight healthy men underwent, in randomized order, an acute bout of high intensity cycling exercise (study day 1) and a hyperinsulinemic-euglycemic clamp (study day 2). Skeletal muscle biopsies were obtained pre and immediately post each intervention (A). Glycogen content (B) and leg glucose balance during clamp (C). Western blot confirmation of insulin and exercise-induced signaling (D). Phosphoproteomics workflow with 2x TMT 11-plex labelling, phosphopeptide enrichment and LC-MS/MS analysis (E). Principal component analysis of insulin and exercise signaling responses (F-G). Volcano plots of insulin and exercise signaling (x-axis = logFC, y-axis = -log10 (*P* value)). A linear model (Limma) was used to test for difference in phosphosite abundance between conditions with a false-discovery rate set to 5% (H-I). *P* < 0.01 = **.

Subsequently, we performed in-depth analysis of the phosphoproteome in insulin and exercise-stimulated skeletal muscle. Proteomic analysis of skeletal muscle is inherently challenging due to the presence of highly abundant contractile proteins hindering the detection of lowly abundant proteins ^32^. To overcome this, we employed stable isotope labelling with 11-plex Tandem Mass Tag (TMT), phosphopeptide enrichment and phosphopeptide-level fractionation prior to tandem mass spectrometry (LC-MS/MS). This approach allowed a comprehensive phosphoproteome in human skeletal muscle and the quantification of 13,717 phosphopeptides (11,934 phosphosites localization probability > 0.75) on 3,368 proteins (Figure 1E). Principal component analysis demonstrated clustering of samples by subject on component 1, and by stimulus on component 2 (Figure 1F-G). Differential abundance analysis of the phosphoproteome revealed significant regulation with 805 and 4,784 sites changing in response to insulin and exercise, respectively (False-discovery rate (FDR) < 0.05, Figure 1H-I). This included known insulin-(upregulation of RPS6 S236/T241, TBC1D4 S341, TSC2 S939 and downregulation of GYS1 S645) and exercise-signaling proteins (upregulation of STIM1 S257, ACACB S222 and downregulation of PDHA1 S300) as well as new insulin signaling proteins, such as the S741 phosphosite on the RNA-binding protein 15, RBM15, which was downregulated in response to insulin. Interestingly, RBM15 was recently associated with liver insulin resistance through epigenetic m6A regulation^33^. Our comprehensive phosphoproteomic analysis encompassing insulin and exercise stimuli in the same individuals represents a valuable resource for the scientific community (Supplementary table 1A-C).

### Shared and Distinct Features of Insulin and Exercise Signaling

To decipher the signaling pathways and regulatory networks activated by insulin and exercise, we investigated if specific kinases were activated and potentially responsible for the observed phosphorylation events in our dataset. Recent work on mapping substrates of the human Ser/Thr kinome has provided *in vitro* evidence of kinase-substrate relationship^34^. Leveraging this work, we overlaid our sequence windows surrounding phosphorylated serine and threonine residues with the *in vitro*-predicted kinase motifs. As substrates are essential for interaction with their upstream kinase, we filtered for kinases known to be expressed in skeletal muscle tissue^35^. Subsequently, we performed a kinase enrichment analysis, predicting activation of known kinases (P70S6K, P90S6K/RSK, SGK1), and a hitherto new insulin-activated kinase, PIM3, in response to insulin (Figure 2A, Supplementary table 1E). Exercise resulted in activation of known kinases such as PKACA, AMPKα1, AMPKα2, MAPKAPK2, MAPKAPK3, RSK2 (Figure 2B, Supplementary table 1F) and new exercise kinases such as QIK (also known as salt-inducible kinase 2, SIK-2). Phosphosites regulated by exercise were enriched for processes related to protein degradation, protein conformation and protein activation/inhibition while those regulated by insulin were associated with intracellular localization (two-sided fishers exact test *P* < 0.05) (Figure 2C). Furthermore, sites with a reported function in molecular associations (i.e., interactions) or disease-associated function were enriched by both insulin and exercise (Figure 2C).

**Figure 2.**
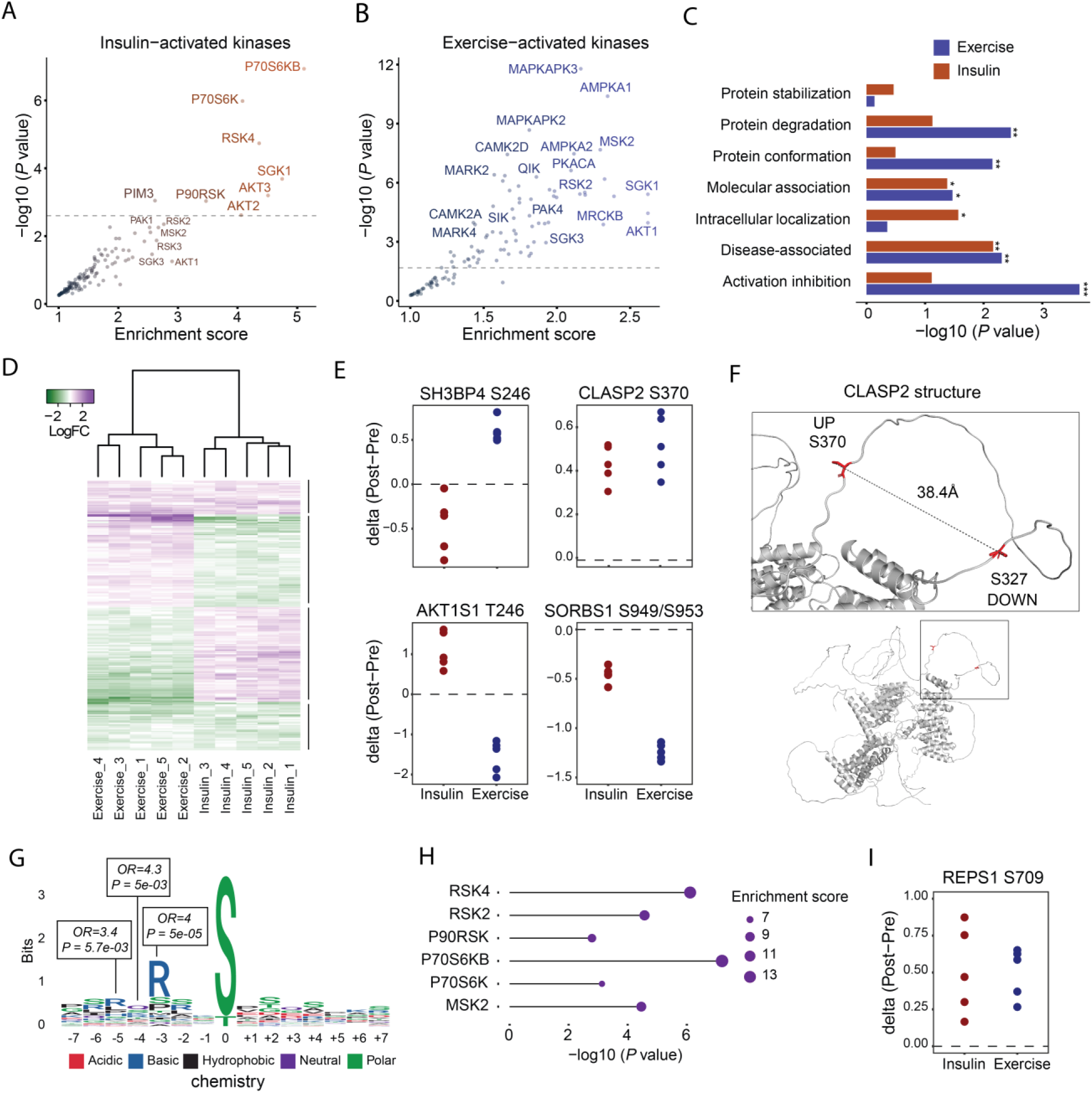
Shared and Distinct Features of Insulin and Exercise Signaling. Predicted insulin- and exercise-activated kinases from the kinase library (A-B). Two-sided Fisher’s exact test of phosphosite functionality based on the PhosphoSitePlus database (C). Heatmap of individual fold-change response of significant phosphosites to exercise and insulin stimulation across the five subjects (D). Selected phosphosites regulated in opposite or the same direction (E). Alpha-fold predicted structure of CLASP2 with highlighted serine residues phosphorylated or dephosphorylated by both insulin and exercise (F). Sequence window of phosphosites upregulated by insulin and exercise. A Two-sided Fisher’s exact test was used to test for residue overrepresentation (G). Kinase enrichment analysis of phosphosites upregulated by both insulin and exercise (H). Individual fold-changes of p-REPS1 S709 in response to insulin and exercise (I). *P* < 0.05 = *, *P* < 0.01 = **, *P* < 0.001 = ***.

To elucidate the crosstalk between insulin and exercise-induced signaling, we filtered for significantly altered sites by both insulin and exercise (< 5% FDR in both comparisons). In total, this analysis revealed 233 phosphosites that were regulated either in the same or opposite direction (Figure 2D & Supplementary table 1C-D). When examining individual responses to exercise and insulin, we observed that the majority of phosphosites, 162 in total, were regulated in opposite directions, compared to 71 sites regulated in the same direction. This highlights the major role of insulin and exercise in regulating muscle anabolic and catabolic metabolism, respectively. One example is the known T246 site of AKT1S1 (also known as PRAS40), which was phosphorylated in response to insulin stimulation and dephosphorylated on the same site in response to exercise (Figure 2E). Conversely, SH3BP4, a protein involved in autophagy and receptor endocytosis, displayed dephosphorylation on S246 in response to insulin and phosphorylation in response to exercise^36^. Exercise and insulin are known to induce and attenuate autophagy in skeletal muscle, respectively^37,38^. Therefore, the S246 phosphorylation on SH3BP4 could be a key regulatory step in skeletal muscle autophagy.

We hypothesized that the phosphosites regulated in the same direction during insulin and exercise, could be involved in metabolic processes such as skeletal muscle glucose uptake. Accordingly, these sites were enriched for biological processes related to exocytosis, actin filament organization and endomembrane system organization (Figure S2A). Similarly, sites downregulated with both stimuli were enriched for cytoskeletal organization (Figure S2B). Exploring specific examples, we observed that SORBS1 (also known as Cbl-associated protein/CAP) was dephosphorylated on both S949/S953 (same phosphopeptide) in response to both insulin and exercise (Figure 2E). SORBS1 is required for insulin-stimulated glucose uptake in adipocytes independent of the canonical PI3K pathway^39^. However, its role in skeletal muscle insulin signaling has been debated. Additionally, we observed that CLIP-associating protein (CLASP)-2 S370 phosphorylation was induced in response to both insulin and exercise (Figure 2E), while CLASP2 S327 was dephosphorylated in response to both stimuli (Figure S2C). When we visualized the AlphaFold predicted structure, we found both sites to be present in the same loop domain suggesting interdependency (Figure 2F). The same observation was seen on the sister-protein CLASP1 at S600 and S559 (Figure S2D-E). CLASP2 is required for insulin-stimulated GLUT4 translocation, thereby serving as a positive readout for our phoshoproteomics analysis^40,41^. In total, we identified 29 phosphosites across 27 proteins induced by both insulin and exercise. To gain further insights into their upstream kinases, we conducted a motif enrichment analysis, revealing a prevalent sequence motif surrounding the phosphosites with a highly significant proportion of arginine at position -3 (Odds Ratio = 4, *P* = 5e-05) (Figure 2G). Additionally, these motifs were enriched for P70S6K and P90S6K (RSKs) kinases (Figure 2H). Interestingly, prior research has shown that S6K and RSK are activated by both insulin and exercise in human skeletal muscle^42–45^. In addition, RSK1 was recently found to induce GLUT4 translocation in adipocytes through TBC1D4 phosphorylation in response to stress stimuli^46^. Of particular interest, we identified the protein RalBP1-associated Eps domain-containing protein 1 (REPS1), to be phosphorylated on S709 in response to insulin and exercise (Figure 2I).

### The Insulin and Exercise-responsive Protein, REPS1, is a Critical Regulator of Skeletal Muscle Glucose Uptake

Given that REPS1 S709 was phosphorylated in response to both insulin and exercise stimuli, we explored its role in skeletal muscle. The amino acid sequence surrounding serine 709 was found to be conserved from zebrafish to human containing the motif favored by AGC kinases^47,48^ (Figure 3A). Moreover, REPS1 has previously been characterized for its involvement in endocytosis^47^. Therefore, we decided to investigate the role of REPS1 in skeletal muscle glucose metabolism. We validated our observation by immunoblotting for phospho-REPS1 at S709 in human skeletal muscle, confirming a significant increase in phosphorylated REPS1 at S709 in response to both insulin and exercise (Figure 3B-C). Strikingly, REPS1 S709 phosphorylation (Post-Pre clamp) tightly correlated with steady-state leg glucose uptake with a squared Pearson’s correlation coefficient of 0.86 (*P* < 0.001) (Figure 3D). In contrast, p-AKT S473 and p-mTOR S2448, classical markers of insulin signaling, showed poor correlation (Figure S3A-B). This suggests a close association between skeletal muscle glucose uptake and REPS1 S709 phosphorylation.

**Figure 3.**
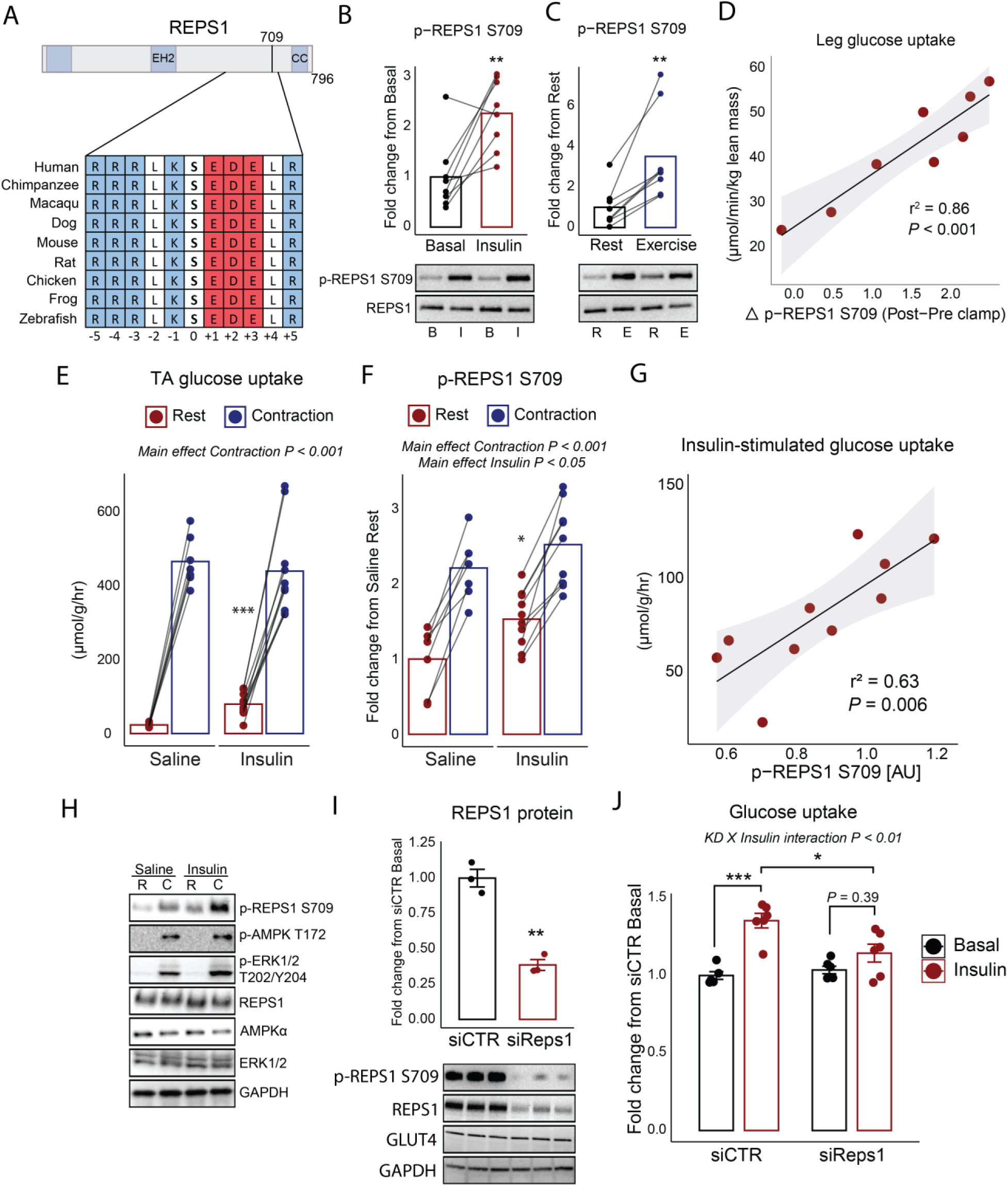
The Insulin and Exercise-responsive Protein, REPS1, is a Critical Regulator of Skeletal Muscle Glucose Uptake. Highly conserved REPS1 sequence window around the serine 709 site (A). Western blot validation of REPS1 phosphorylation in human skeletal muscle before (basal) and after (insulin) a two-hour hyperinsulinemic euglycemic clamp (B) and at rest and after 10 minutes high intensity cycling exercise (C). Pearson’s correlation analysis of steady-state leg glucose uptake and delta (Post-Pre clamp) REPS1 S709 phosphorylation (D). In vivo 3H-2-deoxy-glucose uptake during rest or *in situ* contractions of tibialis anterior (TA) muscle with saline or 15 mU insulin stimulation (E). Pearson’s correlation analysis of insulin-stimulated glucose uptake into TA muscle and REPS1 S709 phosphorylation (F). Representative western blot from *in situ* contraction experiment (H). Western blot validation of SiRNA-mediated knockdown of *Reps1* in C2C12 myotubes (I). Insulin-stimulated 3H-2-deoxy-glucose uptake in C2C12 myotubes transfected with control or *Reps1* siRNA (J). Data in B and C were analyzed with a two-sided paired sample t-test (n = 8). Data in E and F were analyzed by a two-way ANOVA with repeated measures and Sidak multiple comparison test (n = 7-10). Data in I were analyzed by a two-sided two sample t-test. Data in J were analyzed by two-away-ANOVA with Tukey’s multiple comparisons test (n=6) (J). *P* < 0.05 = *, *P* < 0.01 = **, *P* < 0.001 = ***.

To ascertain the conservation of response, we anaesthetized C57BL/6NTac male mice and surgically stimulated one leg to contract while the contralateral leg remained rested. Upon contractions, mice were injected with either saline or 15 mU insulin combined with ^3^H-2-deoxyglucose for 15 minutes. Tibialis Anterior (TA) muscle had a ∼20-fold and 3.4-fold increase in glucose uptake during contractions and insulin, respectively, however no additive effect was observed (Figure 3E). We also measured REPS1 S709 phosphorylation and found increased phosphorylation in all contracted muscles in response to 15 minutes of *in situ* contraction (main effect *P* < 0.001) (Figure 3F & H). As observed in humans, 15 minutes of *in vivo* insulin stimulation increased REPS1 S709 phosphorylation (main effect *P* < 0.05). The insulin-stimulated glucose uptake into TA muscle had a strong correlation with REPS1 S709 phosphorylation (r^2^ = 0.63, *P* = 0.0063) (Figure 3G) whereas contraction-stimulated glucose uptake showed poor correlation (r^2^ = -0.064, *P* = 0.58) (Figure S3C). To test if REPS1 is required for glucose uptake, we transfected C2C12 muscle cells with small interfering RNA (siRNA) against mouse *Reps1*. Efficient knockdown of *Reps1* was validated at the protein level by Western blot analysis (p<0.01), without any significant changes in GLUT4 expression (Figure 3I). Insulin-stimulated ^3^H-2-deoxyglucose was significantly impaired when REPS1 protein expression was reduced (Interaction *P* < 0.01) (Figure 3J). Collectively, these findings establish REPS1 as a critical node for insulin and exercise signaling and hence in the regulation of muscle glucose uptake.

### RSK is an Upstream Kinase of REPS1 S709 and is Associated with Vesicle-Sorting Proteins in Skeletal Muscle

We hypothesized that a single upstream kinase could be activated during insulin and exercise stimulation and be responsible for the increased phosphorylation of REPS1 S709. Interestingly, the sequence motifs of phosphosites upregulated by both insulin and exercise were enriched for the P70S6K and P90S6K isoforms (Figure 2H). In support, a recent study found that RSK can phosphorylate REPS1 at S709 *in vitro* in 293T cells^49^. To investigate if RSK is involved in insulin-stimulated glucose uptake, we preincubated C2C12 myotubes with the RSK inhibitor, BI-D1870, for 20 minutes followed by 10 minutes of maximal insulin stimulation. We observed a significant increase in glucose uptake and REPS1 S709 phosphorylation in response to insulin, however these insulin-induced effects were blunted in cells where RSK was inhibited (interaction *P* = 0.003, *P* = 0.075) (Figure 4A-C). Importantly, the RSK inhibitor lowered phosphorylated REPS1 without affecting insulin-stimulated phosphorylation of AKT on S473. Furthermore, the basal level of phosphorylated REPS1 on S709 was lowered suggesting RSK is the upstream kinase.

**Figure 4.**
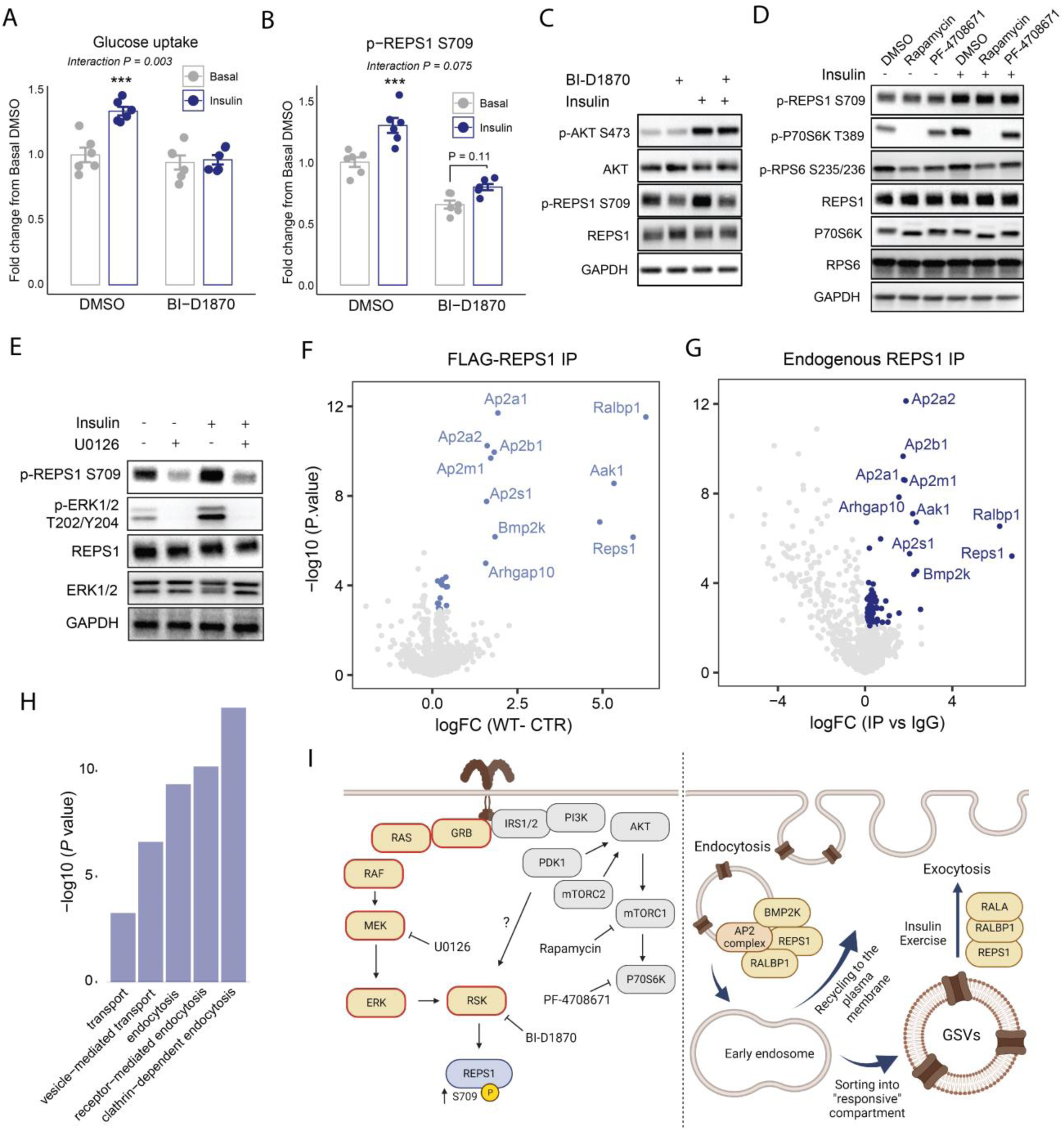
RSK is an Upstream Kinase of REPS1 S709 and is Associated with Vesicle-Sorting Proteins in Skeletal Muscle. Insulin-stimulated (100 nM, 10 minutes) glucose uptake in C2C12 myotubes preincubated (20 minutes) with DMSO or 10 µM of the RSK inhibitor, BI-D1870 (A). Quantified western blot analysis of signaling as in A (B). Representative western blot analysis for signaling in B (C). As in A-C, but preincubated with DMSO, Rapamycin (100 nM) or PF-4708671 (10 µM) for 20 minutes before 10 minutes of 100 nM insulin stimulation (D). As in A-D, but preincubated with 10 µM U0126 for 20 minutes before 10 minutes of 100 nM insulin stimulation (E). Flag-Pull-down of C2C12 myotubes transduced with either FLAG-*Reps1* vs FLAG-CTR (F). Differentially enriched proteins highlighted in blue (two-sample t-test, FDR < 5%). Endogenous IP of REPS1 vs IgG control IP (G). Differentially enriched proteins highlighted in dark blue (two-sample t-test, FDR < 5%) (G). GO-enrichment analysis of significant interactors found in both pull-down and endogenous IP experiments (H). Illustration of signaling pathways affected and the proposed role of REPS1 in vesicle trafficking (I). Illustration made in Biorender. *P* < 0.001 = ***. Data in A and B were analyzed by two-away-ANOVA with Tukey’s multiple comparisons test.

To test whether insulin-induced REPS1 S709 phosphorylation is also dependent on the mTOR-S6K pathway, we stimulated C2C12 myotubes with Rapamycin and PF-4708671, known inhibitors of mTORC1 or P70S6K, respectively. These inhibitors reduced insulin-stimulated phosphorylation of P70S6K T389 (mTOR site) and RPS6 S235/236 (P70S6K site), without affecting REPS1 S709 phosphorylation (Figure 4D). As RSK can also be activated by 3-phosphoinositide-dependent protein kinase 1 (PDK1), we pre-treated cells with a PDK1 inhibitor, MK-7, but found no impairment in insulin-stimulated phosphorylation of REPS1 S709 despite markedly reduced phosphorylation of the PDK1 site, T308 on AKT (Figure S4A). RSK is also known to be activated through the RAF-MEK-ERK pathway^50,51^ (Figure 4I, left). Therefore, we inhibited the MAPK-pathway and observed concurrent reduction in ERK1/2 T202/Y204 and REPS1 S709 phosphorylation indicating that signaling from the insulin receptor to REPS1 S709 is mediated through the MAPK pathway (Figure 4E).

The serine 709 residue of REPS1 lies within a disordered region where phosphorylation could trigger gain/loss of protein-interactions. To identify REPS1 binding partners, we applied adeno-associated virus (AAV) assisted overexpression of FLAG-control or FLAG-*Reps1*. We next performed mass spectrometry-based interaction proteomics. As expected, REPS1 was the most enriched protein in the WT-Flag IP vs CTR-Flag (Figure 4F). RALBP1, a protein recently shown to bind REPS1 and regulate GLUT4-exocytosis in fibroblast, co-precipitated with REPS1 in our screen^52^. Furthermore, all five subunits of the endocytotic adaptor protein 2 (AP-2) complex (Ap2a1, Ap2a2, Ap2b1, Ap2m1 and Ap2s1) and associated kinases, Aak1, Bmp2k, were bound to REPS1. These interactors were similarly found by immunoprecipitation and MS-based proteomics of endogenous REPS1 (Figure 4G). The REPS1 interactors confidently found in both Flag and endogenous immunoprecipitation were enriched for processes related to endocytosis and vesicle transport (Figure 4H & S4B). In summary, we show that REPS1 binds to proteins related to both the exocytosis and endocytosis machinery (Figure 4I, right), and our findings reveal critical upstream and downstream signaling nodes for REPS1, shedding light on its intricate regulation and involvement in skeletal muscle glucose uptake.

### Insulin-induced REPS1 S709 Phosphorylation *in vivo* is Impaired in Multiple Models of Insulin Resistance

After establishing the critical role of REPS1 in skeletal muscle glucose uptake, we sought to investigate whether REPS1 phosphorylation is impaired under insulin resistant conditions. To elucidate if insulin-stimulated phosphorylation of REPS1 is affected by insulin resistance, we fed C57BL/6NTac male mice a low-fat diet (LFD) or high-fat diet (HFD) for 16 weeks, injected them acutely with saline or insulin (1U/kg) and tested their blood glucose levels before and 15 minutes following injection (Figure 5A). Mice fed a HFD had higher body weight (*P* < 0.001) and, upon insulin injection, displayed less reduction in blood glucose compared to LFD-fed mice (interaction *P* < 0.05, Figure 5B), indicating reduced whole-body insulin sensitivity in the HFD group. Intriguingly, the insulin-stimulated phosphorylation of REPS1 at S709 was lower in mice fed a HFD compared to those fed a LFD (interaction *P* < 0.05) (Figure 5C). This effect was specific to skeletal muscle, as epididymal white adipose tissue (eWAT) and liver REPS1 phosphorylation was not affected by insulin or diet. This was despite lowered insulin-stimulated AKT S473 phosphorylation in all three tissues. Furthermore, the insulin-induced drop in blood glucose levels correlated with skeletal muscle REPS1 S709 phosphorylation levels (r^2^ = 0.69, *P* = 0.01) (Figure S5A).

**Figure 5.**
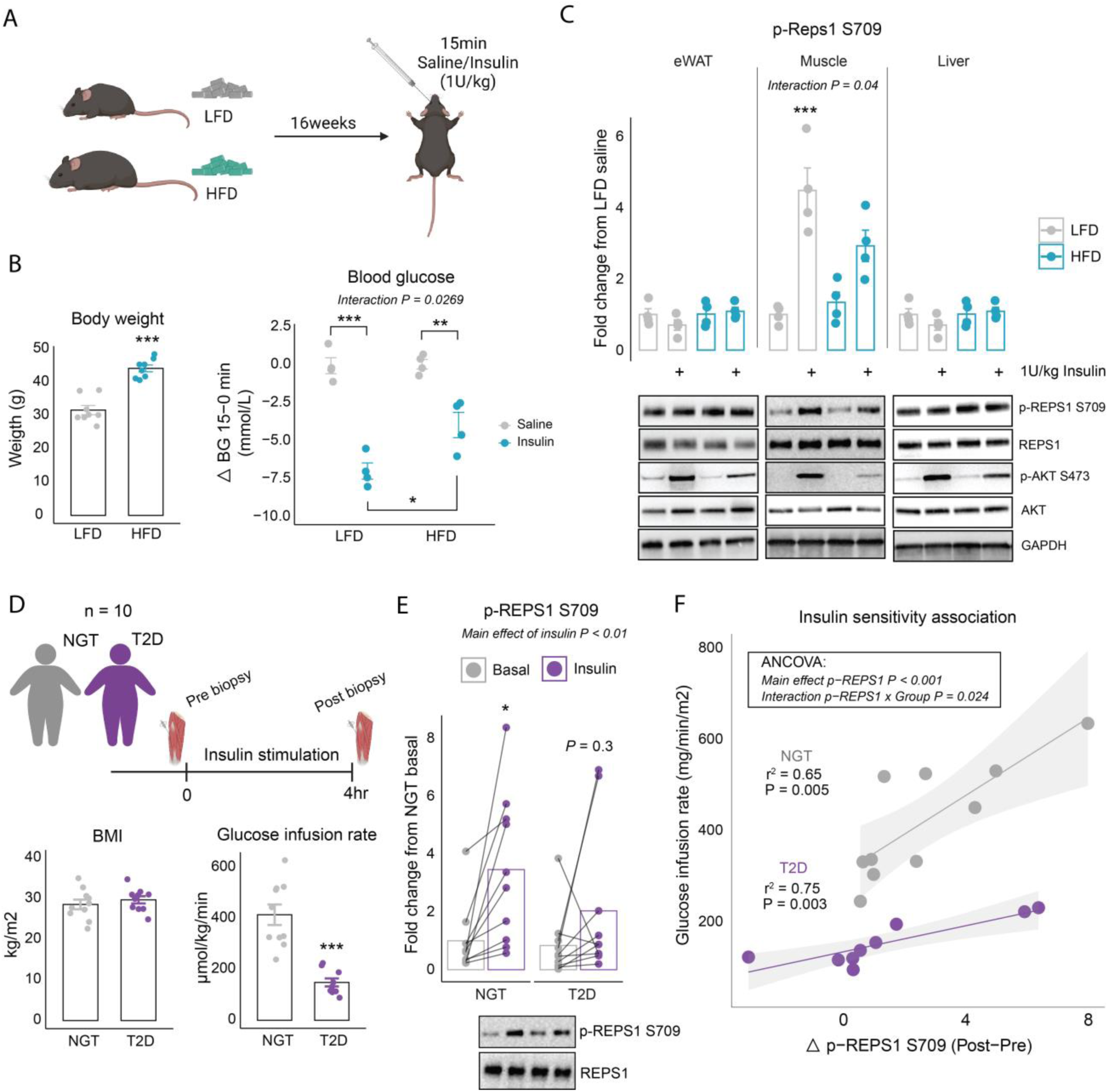
Insulin-induced REPS1 S709 phosphorylation *in vivo* is Impaired in Multiple Models of Insulin Resistance. Sixteen C57BL6 male mice (7 weeks old) were fed a low-fat diet (LFD; n = 8) or high-fat-diet (HFD; n = 8) for 16 weeks. At day of termination, mice were injected with either saline or 1U/kg insulin and tissues were collected after 15 minutes (n = 4) (A). Body weight of mice after 16-weeks of diet intervention and blood glucose (mM) delta (post-pre) values in response to saline/insulin injection (B). Corresponding western blot analysis of insulin signaling in epidydimal white adipose tissue (eWAT), quadriceps muscle and liver tissue from same animals (C). BMI and glucose-infusion rate (GIR) in response to a hyperinsulinemic euglycemic clamp (4 hrs) of 10 normal glucose tolerant (NGT) and 10 patients with T2D (D). Quantified Western blot of REPS1 S709 phosphorylation pre and post insulin stimulation (E). Association of GIR and delta REPS1 S709 phosphorylation in skeletal muscle (F). Data in B-C were analyzed by a two-way ANOVA and Tukey’s multiple comparisons test. Data in E were analyzed by a two-way ANOVA with repeated measures and Sidak multiple comparisons test. Data in F were analyzed by ANCOVA analysis followed by individual Pearson correlation (F). *P* < 0.05 = *, *P* < 0.01 = **, *P* < 0.001 = ***.

To determine if this impairment is present in humans, we analyzed skeletal muscle biopsies from 10 people with T2D and 10 controls matched by age, sex, and BMI with normal glucose tolerance (NGT) in the resting, basal and insulin-stimulated steady-state periods of an hyperinsulinemic euglycemic clamp (Figure 5D). People with T2D exhibited lowered insulin sensitivity than controls (*P* < 0.001) (Figure 5D). REPS1 S709 phosphorylation was significantly increased in response to insulin infusion by 3.5-fold (*P* < 0.05) in the NGT group compared to 2.0-fold in the T2D group (*P* = 0.3) (Figure 5E). We observed a highly heterogeneous response in both groups. Therefore, we correlated the delta (Post-Pre clamp) response in phosphorylated REPS1 S709 with whole-body insulin sensitivity. Strikingly, there was an interaction in responses between individuals with T2D and NGT (ANCOVA interaction *P* = 0.024) (Figure 5F). Therefore, we tested individual associations within each group and found significant correlation for both groups (NGT: r^2^=0.65, *P* = 0.005, T2D: r^2^=0.75, *P* = 0.003) (Figure 5F). Together, these results highlights the role of REPS1 as a key signaling protein impaired in insulin resistance.

### *REPS1* Variants’ Association with Complex Traits, mRNA Expression, and Splicing

After identifying that REPS1 was closely linked to glucose uptake and insulin sensitivity, we explored whether *REPS1* gene variants are associated with cardiometabolic health. By analyzing the NHGRI-EBI Genome-wide association studies (GWAS) catalog, we identified 17 *REPS1* variants linked to 22 distinct phenotypes (*P* < 1×10^-6^)^53–57^. Using the 1000G European reference panel for linkage disequilibrium (LD)-pruning, we discerned 6 independent *REPS1* signals (r^2^ < 0.1). Employing Open Target Genetics, we focused on the lead variant with the most significant associations within each signal (Figure 6, Supplementary Table 2A-B). Remarkably, 4 out of these 6 independent lead variants (rs7870658, rs2750415, rs66883945, and rs62441843) are associated with lipid traits. Two lead variants (rs66883945, rs62441843), located between *REPS1* and *ABRACL*, also share associations with appendicular lean mass, standing height, and sex-hormone binding globulin (SHBG). Intriguingly, two lead variants within *REPS1* introns (rs7870658, rs2750415) are associated with cardiometabolic diseases in type 2 diabetes, such as coronary artery disease and chronic kidney disease. Finally, two other lead variants in *REPS1* introns are singularly associated with corneal-related traits (rs12193050) and leisure screen time (rs200307517), respectively.

**Figure 6.**
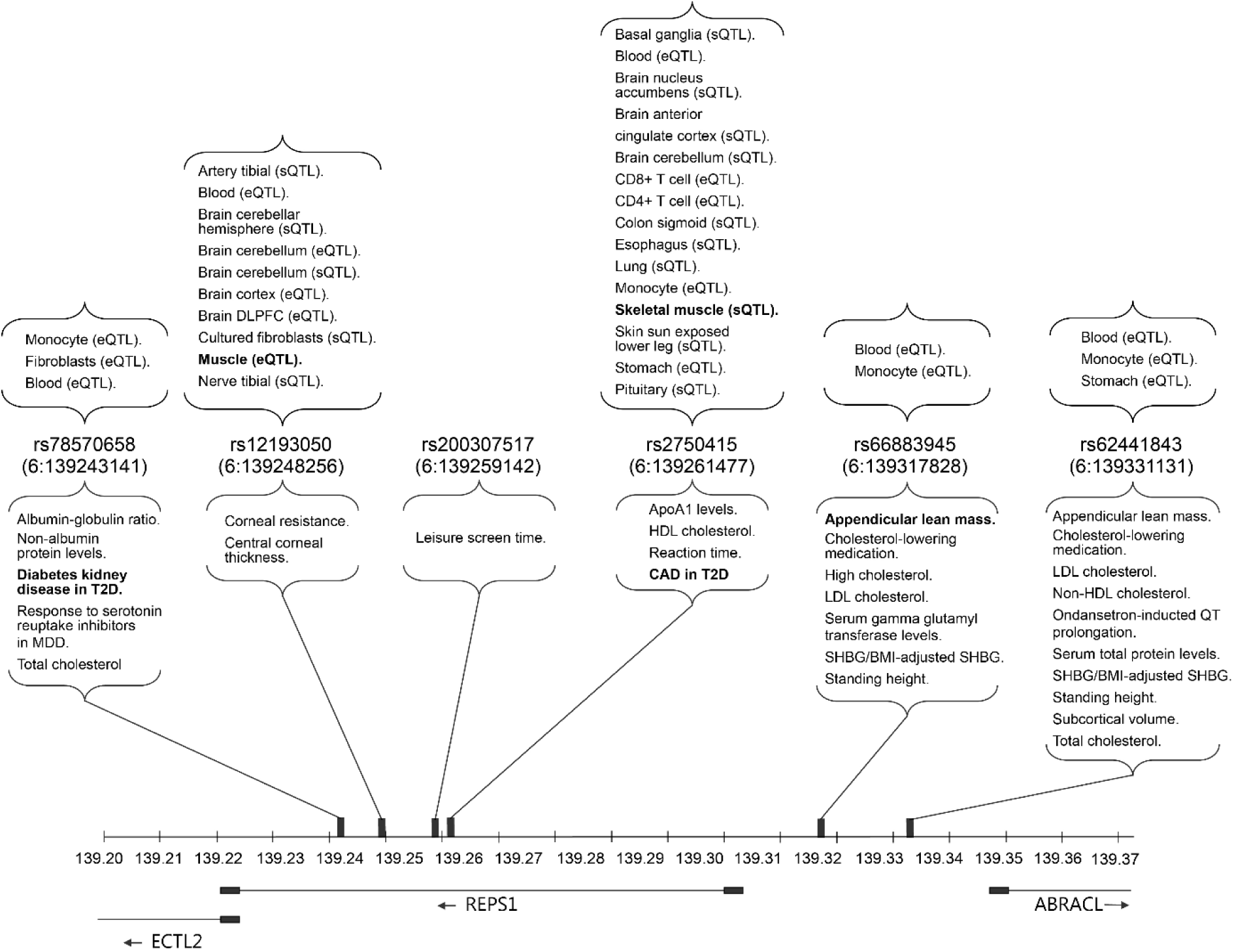
REPS1 Variants’ Associations with Complex Traits, mRNA Expression, and Splicing. Lower segment: Representation of the phenotypes associated with *REPS1* variants using data from NHGRI-EBI GWAS Catalog and Open Target Genetics. In total we identified 6 independent signals (r2 < 0.1) in the *REPS1* region: 4 were located in *REPS1* introns and 2 in an intergenic region between *REPS1* and *ABRACL*. Each signal is represented by their lead variant. Upper segment: Representation of tissues and cell types with significant changes in *REPS1* expression (eQTL) or alternative splicing (sQTL) as reported in Open Target Genetics. We report all tissues and cell types with significant eQTL or sQTL associations for any of the 6 independent *REPS1* lead variants. See Supplementary Tables 2A-C for more information on the associations for each lead variant.

To evaluate the association of the 6 independent *REPS1* lead variants with differential expression (eQTL) and alternative splicing (sQTL) of *REPS1* across tissues and cell types, we queried each variant in Open Target Genetics (Figure 6, Supplementary Table 2C). The majority of the associations were linked to altered expression in blood (5/5 lead variants), monocytes (4/5 variants), brain (2/5 variants) and fibroblasts (1/5 variants). However, two lead variants displayed association to muscle tissue: rs12193050 as an eQTL for muscle, and rs2750415 acts as an sQTL for skeletal muscle. This suggests a potential mechanism in which these genetic variants influence muscle-specific gene regulation of *REPS1*.

## Discussion

Insulin and exercise potently stimulate glucose uptake in skeletal muscle. Here, we mapped the phosphoproteome stimulated by insulin and exercise in human skeletal muscle from the same healthy individuals. We revealed overlapping and unique signaling events triggered by insulin and exercise. REPS1 emerged as a key protein phosphorylated at S709 in response to both exercise and insulin, significantly impacting skeletal muscle glucose uptake. Moreover, phosphorylation of REPS1 strongly correlated with insulin sensitivity and was specifically impaired in skeletal muscle under insulin-resistant conditions, emphasizing its relevance in metabolic diseases such as T2D.

Insulin and exercise induced substantial changes in protein phosphorylation, particularly exercise, which affected 50% of the phosphoproteome. While most of these changes are in line with previous phosphoproteomic studies^29–31^, this study stands out as the first to comprehensively map the deep phosphoproteome in response to acute insulin and exercise stimuli within the same individuals. The study design effectively minimized inter-subject variability and allowed us to precisely pinpoint both distinct and shared signaling events induced by these stimuli. This unique dataset has the potential to serve as a valuable resource for generating novel hypothesis. From our analysis, 71 phosphosites consistently responded to both insulin and exercise, some of which potentially affect glucose uptake. We highlight a fascinating (de)phosphorylation event on the protein CLASP2, where insulin and exercise both induced dephosphorylation of the S327 residue and phosphorylation of the S370 residue, close in space in a disordered region. This mechanism was also observed on the sister protein CLASP1 (Site S559 and S600). CLASP2 is known to regulate insulin-stimulated microtubule dynamics and is directly associated with GLUT4 and required for insulin-stimulated glucose uptake^40,41^. Therefore, CLASP1/2 are likely core proteins required for exercise/insulin-stimulated GLUT4 translocation. Additionally, we also identify vesicle-associated membrane protein-associated protein A (VAPA, also known as VAP-33) to be phosphorylated on S214 by both insulin and exercise. VAPA associates with vesicle-associated membrane protein 2 (VAMP2) and regulates insulin-dependent insertion of GLUT4 into the plasma membrane^58^. Again, this suggests, that insulin and exercise stimulate an overlapping set of proteins important for translocation and insertion of GLUT4-vesicles into the plasma membrane.

Our major discovery centers on the phosphorylation of REPS1 on S709 in response to insulin and exercise and its critical role in skeletal muscle glucose uptake. REPS1 was first characterized as a regulator of endocytosis and epidermal growth factor receptor (EGFR) internalization^47,48^. However, a recent study in HeLa cells found no effect of REPS1 knockout on EGFR internalization, but displayed impaired transferrin receptor (TfR) recycling when REPS1 lacked a phosphorylatable S709 site, emphasizing the functional importance of this site^59^. Considering GLUT4’s presence in TfR-positive vesicles, which are responsive to insulin^6^, we propose that REPS1 has a role in regulating GLUT4 recycling. Our study strongly indicates REPS1 is binding the AP-2 complex and associated kinases (BMP2K, AAK1), suggesting a role in endocytosis, although the specific stage of REPS1 interaction with the endocytic machinery remains unknown. Another recent study found that REPS1 forms a complex with RALBP1 in fibroblasts regulating exocytotic processes including trafficking of GLUT4^52^. As we observed a substantial co-enrichment of RALBP1 with REPS1, we interpret this as strong evidence for a role in GLUT4 exocytosis in skeletal muscle as well. Taken together, we propose a dual role for REPS1: 1) sorting of GLUT4 vesicles in early endosome during endocytosis and 2) acute exocytosis processes possibly though interaction with RALBP1/RALA.

Multiple kinases have been implicated in exercise- and insulin-stimulated glucose uptake in skeletal muscle. Here we identify that phosphosites induced by both insulin and exercise are predicted S6K and RSK kinase family substrates. In addition, acute inhibition of RSK blocked insulin-stimulated glucose uptake in skeletal muscle cells. The role of RSK in glucose uptake is also supported by a recent study that showed a direct interaction and phosphorylation of TBC1D4 with RSK in adipocytes^46^. Here RSK1 was required for stress-stimulated glucose uptake. Interestingly, our findings corroborate a similar axis involving RSK, as we identify RSK as an upstream kinase of REPS1. RSK activation can occur through multiple pathways including the MAPK pathway and by direct phosphorylation by PDK1^60^. In our study, we did not find evidence that PDK1 or the AKT-mTOR-S6K pathway regulate REPS1 S709 phosphorylation in response to insulin stimulation. Instead, we observed REPS1 phosphorylation is regulated via a MEK-ERK-RSK-REPS1 signaling axis, and this study adds REPS1 as a new signaling node downstream of the MAPK-RSK pathway important for skeletal muscle glucose uptake.

A remarkable finding was the strong correlation of human skeletal muscle insulin-stimulated leg glucose uptake and increase in REPS1 S709 phosphorylation. 86% of the variance in leg-glucose uptake could be explained by changes in REPS1 S709 phosphorylation. While this does not establish causality, our study demonstrated impaired insulin-stimulated glucose uptake upon lowering REPS1 expression and S709 phosphorylation in muscle cells strongly suggest a direct role of REPS1 in insulin stimulated muscle glucose uptake. However, from the current study, the role of REPS1 in exercise-mediated glucose uptake is less clear. We did not find an association between REPS1 S709 phosphorylation and contraction-mediated glucose uptake in mice. However, as we suspect the contracted muscle was reaching a maximum limit of glucose transport (supported by no additive effect of contraction and insulin stimulation), further studies with different muscle contraction protocols could shed light on this. However, it is also possible that REPS1 plays a role in the insulin-sensitizing effects of exercise on GLUT4 trafficking potentially through endocytosis/vesicle-sorting of GLUT4 into an insulin-responsive compartment.

Insulin-stimulated glucose uptake into peripheral tissues (e.g., muscle, adipose tissue) or inhibition of glucose production (liver) are compromised in the insulin resistant state. Therefore, we hypothesized that insulin-stimulated phosphorylation of REPS1 at S709 would also be affected in these metabolically active tissues. Despite high expression of REPS1 in both the liver and WAT, insulin-stimulated phosphorylation of REPS1 at S709 was unique to skeletal muscle. This signaling was diminished in the muscles of insulin-resistant mice. Skeletal muscle contains a high expression of GLUT4 and the translocation of GLUT4-containing vesicles to the plasma membrane can be driven by multiple stimuli including muscle contractions^61^. Therefore, it is likely that there exists a unique signaling pathway that governs GLUT4 plasma membrane translocation in skeletal muscle.

Translating our findings in mice to human skeletal muscle, we observed considerable variability in insulin-stimulated REPS1 phosphorylation among individuals with normal glucose tolerance and those with T2D. Interestingly, this variability aligned closely with individual differences in overall insulin sensitivity. This alignment leads us to posit that REPS1 may serve as a pivotal signaling node in humans, holding substantial clinical relevance for metabolic diseases characterized by insulin resistance. In this line, in published GWAS we found that two independent lead variants (rs12193050, rs2750415) were associated with expression and pre-mRNA-splicing in muscle and skeletal muscle tissue, respectively, implicating potential population heterogeneity in REPS1 (isoform) levels. Several genetic variants were related to lipid traits, lean mass, and T2D-related complications. While we anticipated more associations related to glycemic control, the limited methods for evaluating skeletal muscle insulin sensitivity might be a contributing factor. Comprehensive GWAS studies employing hyperinsulinemic clamp techniques are essential, as they could reveal previously undetected associations within genes like *REPS1*, providing a more detailed picture of muscle insulin sensitivity.

In conclusion, we mapped a comprehensive insulin and exercise-induced phosphoproteome in skeletal muscle from healthy individuals. We identified a total of 233 phosphorylation events shared between the two stimuli with 71 residues on 55 proteins regulated in the same direction, and many of which have a role in vesicle trafficking. We thereby provide a comprehensive resource for future studies investigating the role of protein phosphorylation on skeletal muscle glucose uptake and metabolism. We highlight the conserved S709 residue of REPS1 as an insulin and exercise signaling convergence point in mouse and humans and demonstrate a tight association between this phosphorylation site and insulin-stimulated skeletal muscle glucose uptake. REPS1 binding partners and the putative upstream kinase, RSK, suggest a dual role in endocytosis and exocytosis regulation. The muscle-tissue specific insulin response and impairment in insulin resistant mice and humans emphasize the clinical relevance of REPS1 as a potential therapeutic target for diseases characterized with insulin resistance, such as T2D.

## Methods

### Subjects and eligibility criteria

Eight healthy male volunteers took part in the study and were selected from a sub-group of a larger study^62^. Prior to inclusion, subjects met for an assessment of eligibility criteria. The criteria were: healthy males, 18–36 years of age, non-smoker, engaged in physical activity for 2–5 h/week, exhibit maximum oxygen uptake (40–60 ml/kg/min, lean mass 55–65 kg or lean mass index 14–22 kg/m^2^, no usage of beta2-agonists or other prescription medicine, and no allergy towards study medication. Withdrawal criteria included experiencing unacceptable side effects, complications related to the study, or non-compliance with the protocol. The study adhered to the 2013 Helsinki Declaration and was approved by the ethics committee of Copenhagen, Denmark (H-4-2014-002). All subjects provided oral and written informed consent before being included in the study. Body composition was assessed by dual-energy X-ray absorptiometry (Lunar DPX IQ, Version 4.7 E, Lunar Corporation, Madison, WI, USA) (DXA) followed by incremental cycling to exhaustion on a bike ergometer for assessment of VO2max by indirect calorimetry (Monark LC4, Monark Exercise AB, Vansbro, Sweden) as previously described^63^.

### Hyperinsulinemic euglycemic clamp trial

Subjects arrived at the laboratory in a fasting state for assessment of insulin-stimulated whole-body glucose disposal during a hyperinsulinemic-euglycemic clamp. They refrained from vigorous physical activity and alcohol for 48 h and caffeine and nicotine for 24 h. Before the clamp, subjects ingested four to five potassium tablets to prevent hypokalaemia (Kaleorid, 750 mg KCl, Karo Pharma, Sweden) and had a catheter placed in the antecubital vein for infusion of glucose and insulin. In addition, catheters were inserted in the femoral vein and artery and a muscle biopsy was sampled during local anesthesia. The clamp was initiated with a 1 min priming dose of 53.5 pmol/kgbw insulin (100 IU/ml, Novo Nordisk, Copenhagen, Denmark) and continued for 120 min with a constant insulin infusion of 8 pmol/kgbw/min. Femoral venous and artery blood samples were collected before the clamp and every 5 min during the clamp for assessment of glucose concentration. Furthermore, blood samples were collected before and during the clamp for determination of plasma insulin. Glucose was infused during the clamp from a 20% glucose solution (Fresenius Kabi, 200 mg/ml) to maintain euglycaemia at ∼5 mM. At the end of the clamp, another muscle biopsy was sampled.

### Cycle exercise trial

On a separate day in a fasting state, a resting muscle biopsy was obtained under local anaesthesia (2 ml lidocaine without noradrenaline (epinephrine), Xylocaine 20 mg/ml, AstraZeneca, Cambridge, UK). Hereafter, subjects completed 10-min bike ergometer exercise with highest possible effort. Immediately, after the exercise, another biopsy was sampled. Hyperinsulinemic-euglycemic clamp trial and the exercise trials (in same eight individuals) were separated by wash out period of around 1-2 weeks.

### T2D study

A subset of 10 patients with type 2 diabetes from the “specialist supervised individualized multifactorial treatment of new clinically diagnosed type 2 diabetes in general practice (IDA)” study^64^ and 10 persons with normal glucose-tolerance matched on sex, BMI, age and smoking took part in the T2D study. Informed consent was obtained from all subjects before participation. The study was approved by the Regional Scientific Ethical Committees for Southern Denmark (Projekt-ID: S-20120186) and was performed in accordance with the Helsinki Declaration.

All participants had normal results on blood screening tests and ECG. An HEC was performed after an overnight fast. The participants were instructed to refrain from physical activity 48 h prior to test. In patients with T2D, glucose, lipid and blood pressure lowering medication was withdrawn one week prior to the clamp studies. In brief, the clamp consisted of a 2-hour basal period with infusion of a primed constant amount of [3-3H]-tritiated glucose to obtain tracer equilibration. This was followed by a 4-hour insulin-stimulated period using an insulin infusion rate of (40 mU · m-2 min-1) to achieve euglycemia (5,0-5,5 mmol/l) in the insulin-stimulated steady-state period. Skeletal muscle biopsies were obtained before and after the insulin infusion period under local anaesthesia (lidocaine).

### Muscle biopsies

All skeletal muscle biopsies were sampled using a modified Bergström needle technique with suction from the vastus lateralis muscle^65^. After sampling, muscle biopsies were cleaned to remove visible blood, connective tissue, and fat and immediately frozen in liquid nitrogen and stored at −80°C until analysis.

### Sample processing & phosphoproteomics

Freeze-dried muscle was powdered and lysed in 4% sodium dodecyl sulfate (SDS), 100mM Tris pH=8.5 with an Ultra Turrax homogenizer (IKA). Samples were immediately boiled for five minutes, tip-probe sonicated and spun down at 16,000g for 10 minutes. The resultant supernatant was protein precipitated using four times the volume of -20°C acetone overnight. The protein pellets were washed 3 times in acetone, air-dried and resolubilized in 4% sodium deoxycholate (SDC), 100mM Tris pH=8.5 with a bioruptor sonicator (30s on/off, 15 cycles). Protein concentration was determined using Bicinchoninic acid (BCA) assay and 1mg of protein was subsequently reduced and alkylated for 5 minutes at 40° Celsius with 5mM tris-(2-carboxyethyl)phosphine (TCEP) and 40mM 2-chloroacetamide (CAA), respectively. Enzymatic digestion was initiated by adding Trypsin and lysC at a 1:100 enzyme:protein ratio and the mixture was digested overnight at 37°C. Protein digestion was stopped by adding 1% trifluoroacetic acid (TFA) and precipitate was cleared by centrifugation at 20,000g for 10 minutes. Supernatant was desalted on C18 cartridges (Sep-Pak, waters) and eluted with 50% acetonitrile (ACN). Peptides were then vacuum dried and concentration was determined by measuring the absorbance at 280/260 (NanoDrop, Thermo Scientific). 200μg of peptides in 100mM HEPES pH=8-5 were then labeled with 400µg of 11-plex TMT (Thermo, #A34808) for one hour followed by 15 minutes of quenching with 0.27% NH2OH. Peptides pre/post exercise (study day 1) or insulin (study day 2) were labeled on two separate 11-TMT-plexes. A pool of peptides across conditions were placed in the last channel 131C. An aliquot of each sample was tested for near-complete labeling efficiency (>99% labeling).

TMT-peptide pools were then desalted on C18 cartridges and eluted with 50% ACN. Prior to phospho-enrichment, the samples were adjusted to 6% TFA, 3mM KH2PO4, and 50% ACN. TiO₂ -beads (GLsciences) were equilibrated in 80% ACN and 6% TFA. Equilibrated TiO₂ -beads were added in a ratio of 12:1 (bead to peptide) for 15 minutes rotating end-over-end at room temperature. Supernatant was passed through a second and third round of enrichment in an 8:1 and 4:1 bead to peptide ratio, respectively. Beads were combined and washed four times in 60% ACN and 1%TFA. Beads were then transferred in 80% ACN and 0.5% acetic acid on top of 2xC8 layer plugged into a p200 tip. Captured phosphopeptides were then eluted in 80% ACN, 1.25% NH4OH and vacuum dried for 20 minutes. 1% TFA was added to the samples and loaded on to equilibrated three layered Styrene-DivinylBenzene – Reversed Phase Sulfonate (SDB-RPS) StageTips. The phosphopeptides were subsequently washed in 1% TFA in isopropanol followed by two washes with 0.2% TFA. Lastly, phosphopeptides were eluted in 80% ACN and 1.25% NH4OH, vacuum dried and resuspended in 20uL 5% ACN and 0.1% TFA.

### Immunoprecipitation and interactome analysis

Upon C2C12 myoblast differentiation, cells were transduced with 100,000 virus per well for 48 hours before media change. On day of harvesting, cells were starved for three hours and lysed in ice-cold lysis buffer (1% triton-x100, 150mM NaCl, 50mM Tris pH=7.4, 1mM EDTA, 6mM EGTA) supplemented with protease and phosphatase inhibitors (Roche). Cell homogenate was incubated end-over-end for 45 minutes before centrifugation at 16,000G for 10 minutes at 4°C. Protein concentration was determined by the BCA assay. A total of 200µg of protein lysate per sample was used for either FLAG- or endogenous IP. For IP, appropriate antibody was added to lysate and left overnight. The following day protein-G agarose beads were added and samples incubated for two hours. Beads were washed two times in lysis buffer followed by two washes in 150mM NaCl, 50mM Tris pH=7.4. Proteins were then eluted, reduced and alkylated from beads in 2M Urea, 1mM dithiothreitol (DTT) and 0.5µg trypsin for one hour. Supernatant was collected and alkylated with 5mM 2-Iodoacetamide (IAA) and digested overnight at 37°C. The following day, digestion was stopped by adding 1% TFA and 20% of eluate was loaded onto equilibrated Evotips ready for measurement.

### LC-MS measurements for phosphoproteome analysis

Desalted and TMT11plex-labeled peptides were resuspended in buffer A* (2% ACN, 0.1% TFA), and fractionated into 16 fractions by high-pH fractionation. In short, 4µg of peptides were loaded onto a 30 cm x 0.25mm Pepsep 1.9um ReproSil C18 120 Å column via an EASY-nLC 1200 HPLC (Thermo Fisher Scientific) in Spider buffer A (1% tetraethylammonium fluoride (TEAF), 0.1% tetraethylammonium (TEA)). Peptide separation was done with a non-linear gradient of 5 – 44 % Spider buffer B (1% TEAF, 0.1% TEA, 80% ACN) at a flow rate of 1.5 µL / min over 62 min. Collection of fractions was done at 60 s interval with a concatenation workflow to achieve a total of 16 fractions. All fractions were then evaporated, and peptides resuspended in buffer A*, and measured using an EASY-nLC 1200 HPLC system (Thermo Fisher Scientific) coupled through a nano-electrospray source to a Tribrid Eclipse mass spectrometer (Thermo Fisher Scientific). Peptides were loaded in buffer A (0.1 % formic acid (FA)) and separated on a 25 cm column Aurora Gen1 (kept at 50° C), 1.6uM C18 stationary phase (IonOpticks) with a non-linear gradient of 5 – 44% buffer B (0.1% FA, 80% ACN) at a flow rate of 400 nL/min over 91 min with a total gradient time of 101 min including washing. Spray voltage was set to 2400 V. Data acquisition switched between a full scan (120K resolution, 50ms max. injection time, AGC target 100 %) and 3 seconds cycle time-controlled data-dependent MS/MS scans (50K resolution, 120ms max. injection time, AGC target 200 % and HCD activation type). Isolation window and normalized collision energy were set to 0.7 and 35, respectively. Precursors were filtered by charge state of 2-6 and multiple sequencing of peptides was minimized by excluding the selected peptide candidates for 45 s. The PrecursorFit filter, Fit error 70% in a window of 1.2Da, was also activated.

### LCMS measurements for interactome analysis

For the interactome experiment, the peptides were subjected to separation using a 15 cm column with an internal diameter of 150 μM, which was packed with 1.9 μm C18 beads (Pepsep brand). This separation process was carried out on an Evosep ONE HPLC system, employing the specific protocol designed for ’30 samples per day’, and peptides were introduced into the system via a CaptiveSpray source equipped with a 20 μm emitter. The mass spectrometric analysis was performed using a timsTOF SCP mass spectrometer from Bruker, functioning in the parallel accumulation-serial fragmentation (PASEF) mode^66^. To elaborate, the DDA-PASEF scan for both MS and MS/MS was configured to encompass a range from 100 to 1700 m/z. The TIMS mobility spectrum was set within the bounds of 0.6 to 1.6 (V cm−2). Both TIMS ramp and accumulation times were precisely fixed at 100 ms. During each cycle, which lasted a total of 1.17 seconds, 10 PASEF ramps were recorded. The MS/MS target intensity was defined at 20,000, with an intensity threshold at 1,000. Furthermore, a stringent exclusion list was applied, set at 0.4 min for precursors within a narrow window of 0.015 m/z and 0.015 V cm−2.

### Data analysis

Raw files were quantified in MaxQuant v2.0.1.0 against the reviewed human fasta (09082021)^67^. Default settings were applied including a minimum peptide length of seven amino acids. Deamidation (NQ), methionine-oxidation (M) and N-terminal acetylation were set as fixed modifications. Phospho (STY) was set as a variable modification. For interactome analysis, raw files were processed in Fragpipe (v.20.0) with default settings and MaxLFQ enabled^68^.

### Bioinformatics analysis

Maxquant output STY-file was imported into Perseus (2.0.7.0)^69^ and phosphopeptides expanded to represent phosphosites. Data was then imported into R studio v4.2.1 and log2-transformed. Data were median-scaled and unwanted variation was removed by normalizing to stably phosphorylated sites with the phosR package^70^. To identify differentially regulated phosphosites, we applied limma with ebayes data smoothening^71^. The linear model was blocked for subjects. *P* values were adjusted with the Benjamini Hochberg (BH) method and adjusted *P* values < 0.05 were considered statistically significant.

Kinase-activation analysis for Ser/Thr kinase was performed by the online tool: https://kinase-library.phosphosite.org/site^34^. Sequence windows were uploaded with phosphorylated residue centralized. Kinases were then filtered to be expressed in human skeletal muscle as part of the human protein atlas database (∼13000 transcripts)^35^. Regulatory- and disease-associated sites were downloaded from PhosphositesPlus^72^. For each subcategory a two-sided fishers exact test was applied.

### Go enrichment analysis

Gene-ontology enrichment analysis was done with the clusterProfiler package^73^. Enrichment was done on the gene-level. Whole phosphoproteome and interactome data was used as background. BH adjusted *P* values < 0.05 were considered significant.

### Alphafold-structure

Predicted 3-dimensional structure of CLASP2 was extracted from alphafold and specific phosphorylated residues were edited in the Pymol software^74^.

### Interactome data analysis

The combined_protein output file was imported into R studio. Data were log2-transformed and filtered for quantification in at least 50% (3 samples) within a group. Missing values were imputed from a normal distribution with a mean downshift of 1.8 and standard deviation of 0.3. A two-sided t-test was used to assess differences between groups. BH adjusted *P* values < 0.05 were considered significant.

### Querying genetic associations of REPS1 variants

To identify which traits are associated with the *REPS1* gene, we searched for associations between *REPS1* variants and complex traits and diseases in genome-wide association studies (GWAS) by querying “*REPS1*” in the NHGRI-EBI GWAS catalogue^53^ on October 2, 2023. To assess whether the reported associations belong to independent signals, we computed a linkage disequilibrium (LD) matrix using the LDLinkR package (v1.4.2)^54,55^. We considered that variants represent independent signals if the r^2^ between the variants was <0.1 within ±500kb. For each independent signal, we identified the lead variant with the lowest *P* value and queried it in Open Target Genetics^57^ to obtain additional associations from a different source (*P* < 1×10^-6^). We also assessed whether these variants were significantly associated with the mRNA expression (eQTL) or splicing (sQTL) of *REPS1* across different tissues by querying Open Target Genetics using otargen R package (v.1.0.0)^75^.

### Animal experiments

All animal experiments were approved by the Danish Animal Experiments Inspectorate (License no. 2017-15-0201-01276 & 2018-15-0201-01493). 7-week old C57BL/6NTac male mice were fed *ad libitum* a LFD (Research diet:D12450Ji) or HFD diet (Research Diet:D12451i) for 16 weeks on a normal 12:12-h light dark cycle. On the day of termination, fed mice were anaesthetized by an intra-peritoneal injection of pentobarbital (10mg/100mL) and injected retro-orbitally with either saline or human insulin (1U/kg). After 15 minutes, tissues were immediately removed and frozen in liquid nitrogen until further analysis. Blood glucose levels were measured pre/post saline/insulin injection by a glucometer.

### In situ contractions

Mice (male, C57BL/6NTac; 12-16 weeks of age) were anesthetized with pentobarbital (2.5 mg/mouse) and kept warm on a heating mat throughout the experiment. The sciatic nerve was exposed and stimulated (0.5 s train every 1.5s at 5V, 0.1ms, 100hz) to contract the lower leg muscles. The contralateral leg to the contracted leg was sham operated and serves as a non-contracted control. A retro-orbital injection was administered immediately prior to nerve stimulation (15 mU insulin or saline, 3 mg glucose, 20 uCi [3H]2-deoxyglucose in gelofusine) and tissues were dissected 15 minutes after glucose tracer, insulin/saline injection and the initiation of contraction. The TA muscle was quickly dissected and snap frozen. Muscles were processed as in^76^. Protein concentration was determined by the BCA-assay (Thermo #23225). Muscle protein lysate was mix with either 0.3M Ba(OH)_2_ and ZnSO_4_ or 4.5% HClO_4_ before centrifugation at 10000 RPM for 5 minutes at room temperature. Supernatant was collected and added to scintillation vials with Ultima Gold scintillation fluid (Perkin Elmer #6013329) before counted on a Hidex 300 SL scintillation counter. Blood was collected every 5 minutes and specific activity calculated as in^76^.

### Cell culture

C2C12 cells were cultured in Dulbeccós Modified Eagle Medium (DMEM #41965-039) supplemented with 10% Fetal Bovine Serum and 1% penicillin streptomycin (P/S). Cells were maintained in an incubator with 37°C and 5% CO_2_. C2C12 myoblasts were grown to ∼80% confluence before initiating differentiation with DMEM supplemented with 2% Horse serum and 1% P/S.

Signaling experiments: C2C12 myotubes were serum-starved for 4 hours before pre-incubating cells with inhibitors (10µM BI-D1870, 10µM U0126, 100nM Rapamycin, 10µM PF-4708671 and 10µM MP-7) for 20 minutes followed by 10 minutes stimulation of 100nM insulin. Cells were then washed twice in ice-cold phosphate buffered saline (PBS) and lysed in 2% SDS, 50mM Tris pH = 7.4. Lysate was then boiled for 5 minutes at 95°C. A small aliquot was used for DC assay protein determination. Sample concentration was adjusted with SDS-buffer and 4x sample buffer (10% 2-mercaptoethanol, 13.3% glycerol and 444mM Tris-HCl pH=6.8).

SiRNA-mediated knockdown: On the same day as differentiation was initiated, cells were transfected with siRNA against *Reps1* (Dharmacon #L-040836-01-0005) or scramble siRNA (Dharmacon #D-001810-10-20). Complexes of Transit-X2 (Mirus #MIR6000), Opti-MEM and siRNA were prepared in a concentration of 25nM and incubated at room temperature for 20 minutes before the siRNA complexes were added to the cells. The culture media was changed every ∼48 hours. Knockdown of *Reps1* was later confirmed by Western Blot.

Aav transduction: Mouse *Reps1* (NM_009048.2, VectorBuilder) with N-terminal FLAG tag was subcloned into Aav serotype 1 vector under a CMV promoter. The control-FLAG contained a random open-reading frame stuffer similarly under a CMV promoter. Vectors were added at a concentration of 10×10^5^ per well in a 6-well plate at day of differentiation induction. The culture media was changed every ∼48 hours.

### Glucose uptake assay

Glucose uptake of C2C12 myotubes was measured with a ^3^H-2-Deoxyglucose (2DG) assay on day 6 of differentiation. Cells were serum starved for 4 hours in serum free DMEM with 1% P/S prior to the assay. After serum starvation cells were washed one time in pre-warmed Krebs Ringer HEPES buffer (140 mM NaCl, 4.7 mM KCl, 2.5 mM, 1.25 mM, 1.2 mM HEPES, 0.2% bovine serum albumin (BSA), pH=7.4) before incubation with KRHepes or 100 nM insulin for 30 minutes. Thereafter, cells were incubated for 5 minutes with ^3^H-2DG (0.4µCi/mL) and 50µM 2DG. Glucose uptake was stopped by washing cells with ice cold KRHepes buffer while the plate was on ice. Cells were then lysed in 2 % SDS and boiled for 5 minutes at 95°C. 100 µL cell lysate was added to 3 mL of Ultima Gold scintillation liquid (Perkin Elmer #6013329). Samples were kept in the dark overnight and ^3^H-2DG uptake was measured the following day on a Hidex 300 SL scintillation counter. ^3^H-2DG uptake was subsequently adjusted for protein content using a detergent compatible (DC)-assay (Bio-Rad). The glucose-uptake experiment was repeated on three separate days with same observed effect. For the RSK-inhibitor glucose uptake experiment, cells were preincubated with 10 µM BI-D1870 for 20 minutes before 10 minutes of 100nM insulin stimulation.

### Immunoblot analysis

#### WB of animal LFD/HFD tissues

Tissues were powdered and lysed in 4% SDS, 100mM Tris pH=7.4 with an Ultra Turrax homogenizer (IKA) and immediately boiled. Samples were tip-probe sonicated and centrifuged at 16,000g for 10 minutes. Supernatant was collected and protein concentration determined by the DC-assay. 4x Sample-buffer (10% 2-mercaptoethanol, 13.3% glycerol and 444mM Tris-HCl pH=6.8) was added to lysate and boiled again for 5 minutes at 95°C.

#### WB of human samples

Powdered muscle was lysed in ice-cold homogenization buffer (10% glycerol, 20 mM Na-pyrophosphate, 150 mM NaCl, 50 mM HEPES (pH 7.5), 1% Igepal, 20 mM β-glycerophosphate, 2 mM Na3VO4, 10 mM NaF, 2 mM phenylmethylsulfonyl fluoride (PMSF), 1 mM ethylenediaminetetraacetic acid (EDTA) (pH 8), 1 mM ethylene glycol-bis(β-aminoethyl)-N,N,N,N-tetraacetic acid (EGTA) (pH 8), 10 µg⋅ ml–1 Aprotinin, 10 µg⋅ ml–1 Leupeptin and 3 mM Benzamidine) with steel beads at 28.5 Hz for 1 min (Qiagen TissueLyser II, Retsch GmbH, Haan, Germany). Homogenate was then incubated for 1 hour end-over-end at 4°C followed by centrifugation for 20 minutes at 18,320g and 4°C. Protein concentration was determined by the BCA assay.

15µg was loaded on a 4-20% acrylamide gradient gel with 18 or 26-wells (Bio-Rad #5671094 & Bio-Rad #5671095) and separated for 20min at 100v followed by 150V for 1 hour. Proteins were transferred with a semi-dry transfer system (Bio-Rad) onto a polyvinedylene difluoride (PVDF)-equilibrated membrane (Bio-Rad turbo-transfer #1704157) for 7 min at 25V. Membranes were blocked in either 2% milk or 3% BSA in Tris-buffered saline (TBS, pH7.4) supplemented with 0.05% tween-20 for 1 hour and incubated overnight with primary antibody on a rocking platform. The following day, membranes were washed 3 times in TBS-T and incubated for 45 minutes with secondary antibody (Bio-Rad: Goat Anti-rabbit #1706515 or Goat-anti mouse #1706516). Lastly, membranes were washed 3x every 10 minutes for a total of 3 times before ECL (Millipore #WBLUF0500) was added and imaged by chemidoc+ system (Bio-Rad). Immunoblots were analyzed in ImageLab software.

#### Antibodies

REPS1 (CST #6404), p-REPS S709 (CST #12005), GAPDH (CST #2118), p-CREB S133 (CST #9198), p-AKT T308 (CST #9275), p-AKT S473 (CST #9271), AKT2 (CST #3063), pan-AKT total (CST #9272), p-ERK1/2 T202/Y204 (CST #9101), ERK1/2 (CST #4696), mTOR (CST #2972), p-mTOR S2448 (CST #2971), p-AMPK T172(CST #2531), AMPKα2 (CST #2532), ACC (CST #3676), p-P70S6K T389 (CST# 9205), P70S6K (CST #2708), p-RPS6 S235/236 (CST #2211), RPS6 (CST #2217), ACTIN (sigma A2066), p-ACC S221 (Millipore 07-303), GLUT4 (Thermo PA1-1065), FLAG (sigma, F7425).

#### Statistical analysis

For analysis of paired human samples a two-sided paired samples t-test was used. For two-factor designs a two-way ANOVA analysis was used with or without repeated measures. Tukeys or Sidak multiple comparison test were used as post hoc test dependent on repeated measures. For unpaired two condition design a two-sided two sample t-test was used. Pearson’s correlation was used to analyze association between two independent variables (normality tested with the Shapiro-Wilk test *P* > 0.05). In figure 5F, an ANCOVA model was used to predict GIR (dependent variable) by Group (NGT/T2D) and delta (post-pre clamp) p-REPS1 levels (independent variables). Additionally, the model included an interaction term to examine the potential differential effect of p-REPS1 levels on GIR across the two groups (NGT/T2D). *P* values < 0.05 were considered significant.

## Acknowledgement

Mass spectrometry analyses were performed by the Proteomics Research Infrastructure (PRI) at the University of Copenhagen (UCPH), supported by the Novo Nordisk Foundation (NNF) (grant agreement number NNF19SA0059305). This work is supported by an unconditional donation from the Novo Nordisk Foundation (NNF) to NNF Center for Basic Metabolic Research (http://www.cbmr.ku.dk (1 July 2018) (Grant number NNF18CC0034900). The T2D study was supported by grants from the Region of Southern Denmark, the Novo Nordisk Foundation (Grant number NNF15OC0015986), and from The Sawmill Owner Jeppe Juhl and wife Ovita Juhl Memorial Foundation. Mario Garcia Ureña and Tuomas O. Kilpeläinen were supported by the NNF grant number NNF22OC0074128.

## Contributions

Conceptualization, MH, JKL and ASD; Human clinical studies, JVS, KH, MH, JB, KE, SJ, AKL, MT; Investigation, JKL, CBL, MGU, AME, FS, JP, JHS, LS, MT, AGF, TOK, JTT; Writing – Original Draft, JKL and ASD; Writing – Review & Editing, All; Funding Acquisition, ASD, MH and KH; Resources, ASD and MH; Supervision, ASD and MH.

## Supplemental Figure Legends

**Figure S1.**
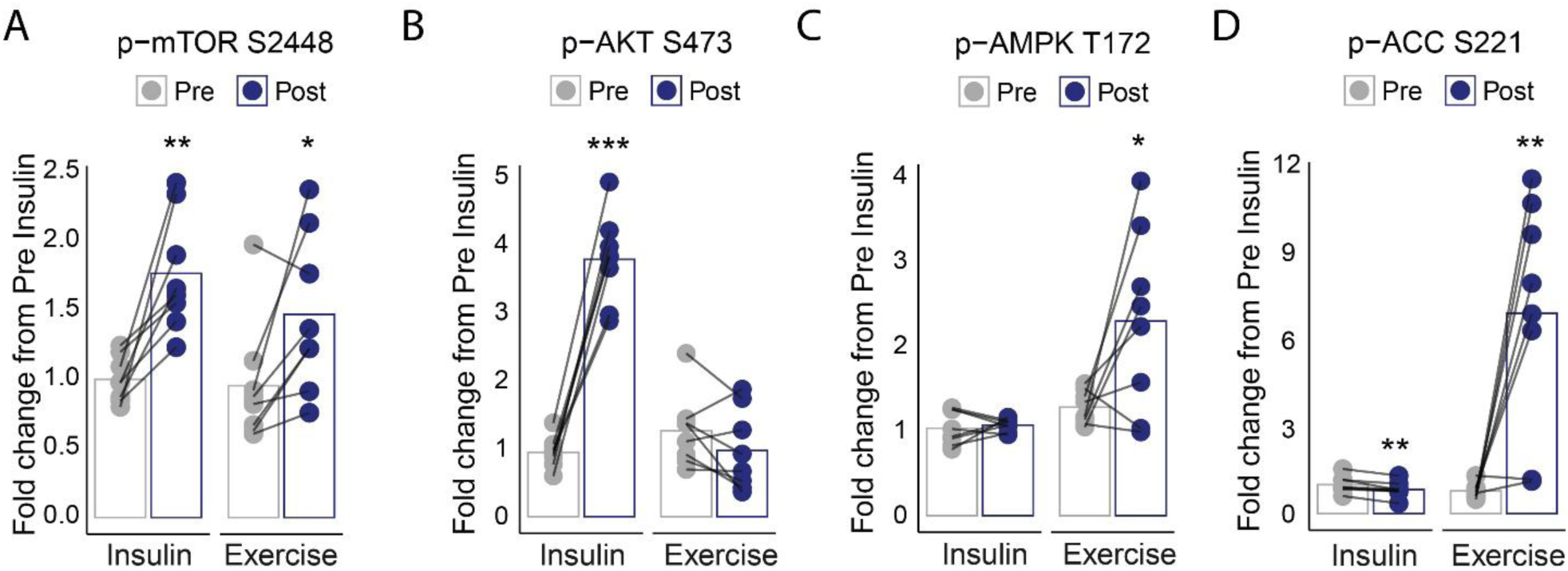
Phosphoproteomic Signature of Insulin and Exercise Signaling in Human Skeletal Muscle. Quantified western blot analysis of p-mTOR S2448 (A), p-AKT S473 (B), p-AMPK T172 (C) and p-ACC S221 (D) in human skeletal muscle pre/post an hyperinsulinemic euglycemic clamp (2hr) and pre/post 10 minutes of high intensity cycling exercise. Paired-Samples t-test was used to analyze mean differences within each experiment. *P* < 0.05 = *, *P* < 0.01 = **, *P* < 0.001 = ***.

**Figure S2.**
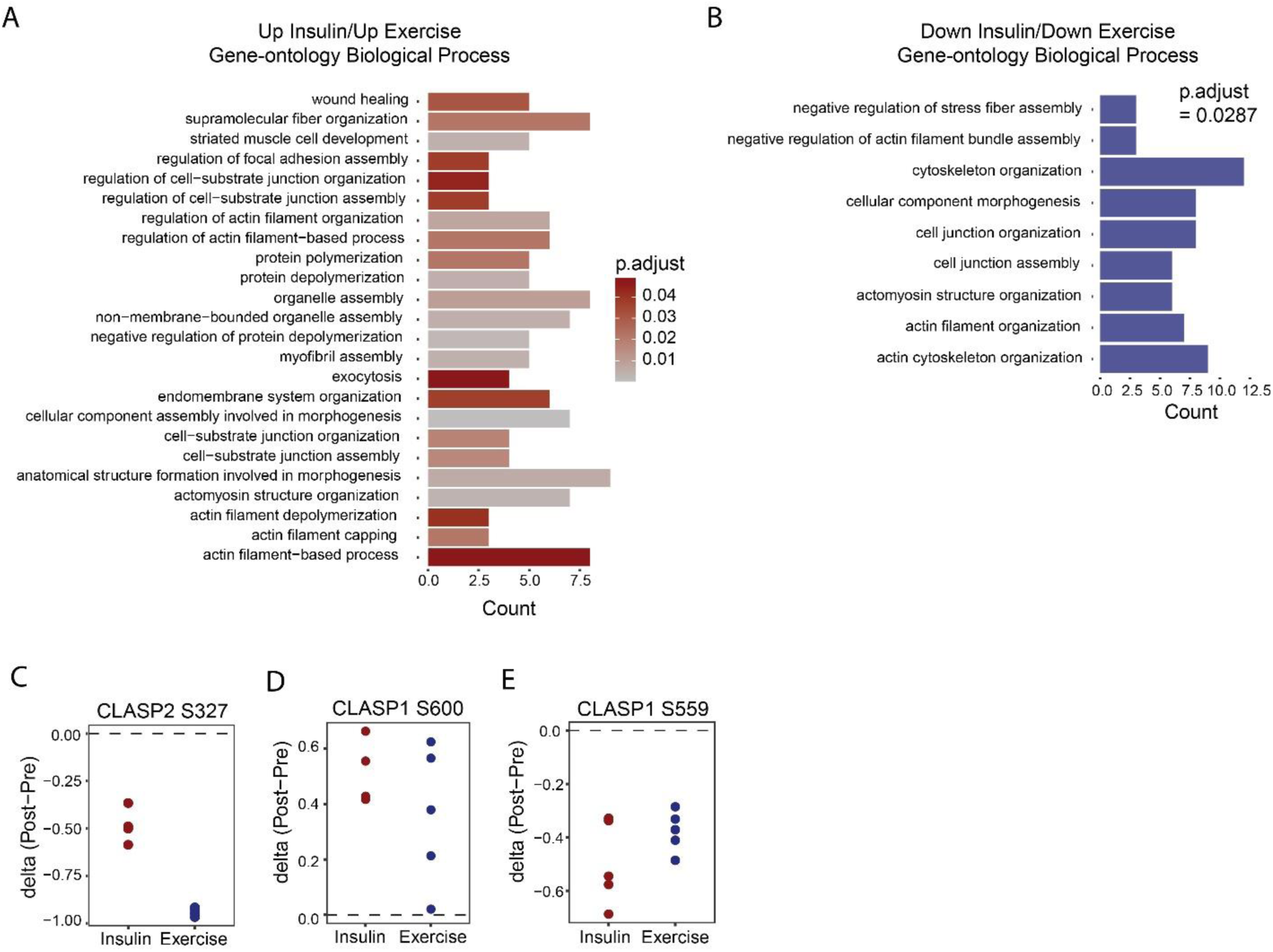
Shared and Distinct Features of Insulin and Exercise Signaling. Gene Ontology (GO) overrepresentation test was used to assess enrichment for Biological Processes within phosphoproteins being phosphorylated (A) or dephosphorylated (B) by both insulin/exercise. The whole phosphoprotein proteome was used as background. Only BP-terms with a Benjamini Hochberg-adjusted *P* value below 5% is shown. Representation of individual fold changes for CLASP2 S327 (C), CLASP1 S600 (D) and CLASP1 S559 (E) in response to insulin stimulation and an acute bout of exercise by LC-MS/MS.

**Figure S3.**
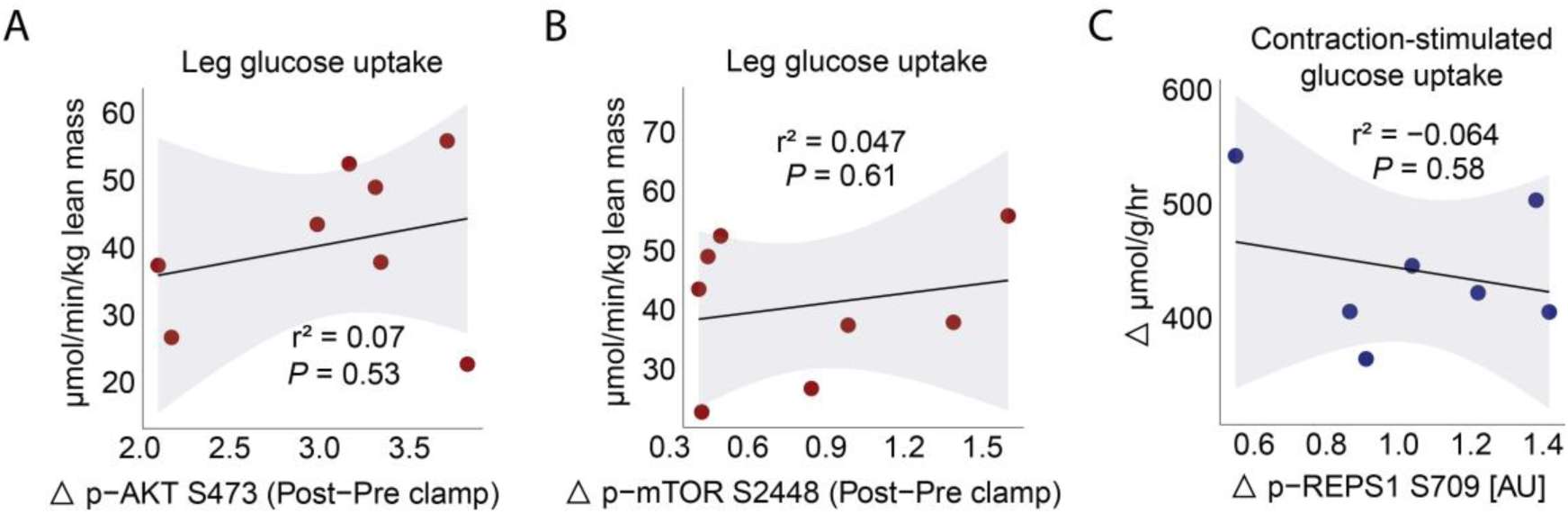
The Insulin and Exercise-responsive Protein, REPS1, is a Critical Regulator of Skeletal Muscle Glucose Uptake. Pearson’s correlation analysis of leg steady-state glucose uptake with delta (Post-Pre) p-AKT S473 (A) and p-mTOR S2448 levels (B) measured by western blot. Pearson’s correlation analysis of delta (Contracted-Rested) glucose uptake and REPS1 S709 phosphorylation (C).

**Figure S4.**
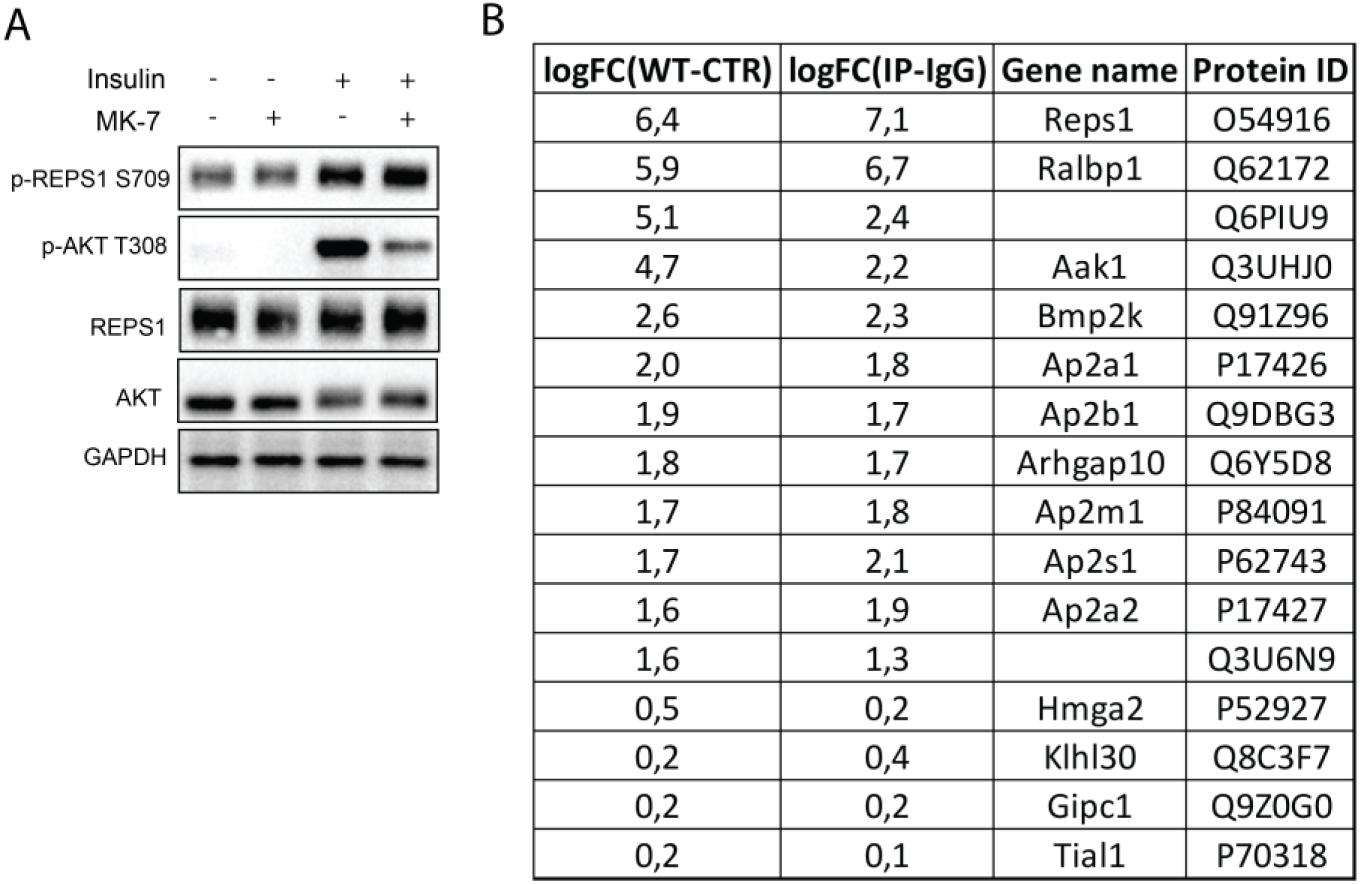
RSK is an Upstream Kinase of REPS1 S709 and is Associated with Vesicle-Sorting Proteins in Skeletal Muscle. C2C12 myotubes were pretreated with 10 µM MP-7 for 20 minutes followed by 10 minutes of 100 nM insulin stimulation. Representative WB is shown in (A). Table showing significant interactors (< 5% FDR) in Flag- and endogenous-IP experiments with log2 fold changes (B). Whole interactome was used as background.

**Figure S5.**
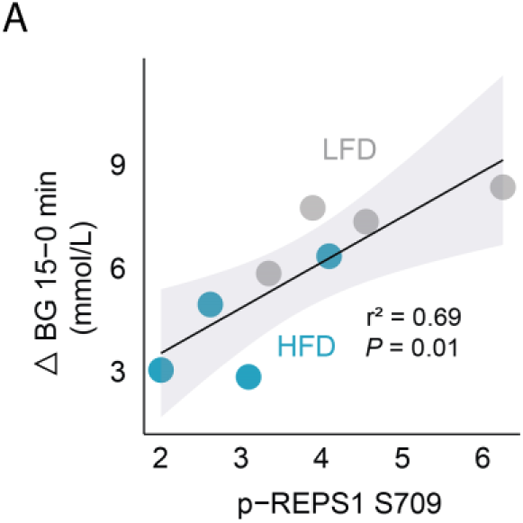
Insulin-induced REPS1 S709 phosphorylation *in vivo* is Impaired in Multiple Models of Insulin Resistance. Association of insulin-lowering (delta 15-0 min) blood glucose (BG) and REPS1 S709 phosphorylation in insulin-stimulated quadriceps skeletal muscle from LFD and HFD fed mice (A).

